# Multi-Region Brain Stimulation Optimization Using Transcranial Direct Current Stimulation

**DOI:** 10.1101/771345

**Authors:** Ziliang Xu, Jinbo Sun, Yao Chen, Yang Yu, Xuejuan Yang, Peng Liu, Badong Chen, Wei Qin

**Affiliations:** Engineering Research Center of Molecular and Neuro Imaging of the Ministry of Education, School of Life Science and Technology, Xidian University, Xi’an, China; School of Electronic and Information Engineering, Xi’an Jiaotong University, Xi’an, China

## Abstract

Transcranial direct current stimulation (tDCS) is a type of noninvasive transcranial electrical brain stimulation. By optimizing the current distribution of each electrode on the scalp, the stimulation can be guided to a target brain region using a tDCS dense electrode array system. However, previous studies have yielded simple results using optimization schemes in single target stimulation cases. The detailed parameter settings for each optimization scheme and the associated simulation results have not been comprehensively assessed. In this study, we investigated parameter settings of optimization schemes in detail in both single target and multi-target cases. Two optimization schemes, minimum least squares (MLS) and maximum electrical field strength (ME), were examined in this study. MLS minimizes the squared errors between the expected electrical field and the estimated electrical field, whereas ME maximizes the electrical field strength in the target region. We constructed a five layer finite-element head model with 64 electrodes placed on the scalp according to the EEG 10/10 system for simulation. We evaluated the effects of stimulation using these two schemes under three conditions, 1) single target stimulation, 2) multi-target stimulation, and 3) multi-target stimulation under specific task activation, which shown that directly using MLS and ME scheme in multi-target stimulation case may lead to a wrong result. We also reported the improved results fixed by our proposed weighted MLS and weighted ME schemes which take detailed parameter settings into consideration. Our results indicate that the parameter settings in each optimization scheme greatly affected the final stimulation results, especially in the case of multi-target stimulation, and thus, indicate that the parameter settings of each optimization scheme should be carefully considered according to the expected stimulation mode. Our results also suggest that, by calculating the parameters through our proposed methods, the weighted ME and weighted MLS scheme can precisely distribute energy into each target brain region.

## Introduction

Transcranial direct current stimulation (tDCS) is a noninvasive brain transcranial electrical stimulation method that has been widely used for many clinical and scientific applications, such as epilepsy [1,2], Parkinson’s disease [3,4], Alzheimer’s disease [5,6], depression [7,8], schizophrenia [9,10] and neuroscience research [11]. tDCS can be used to induce long-lasting alterations of cortical excitability and activity by continuously applying a constant low intensity current through scalp electrodes for a sufficient duration of time [12,13]. Unlike transcranial magnetic stimulation, which generates supra-threshold activation of brain neurons via short-lasting high intensity electromagnetic currents, tDCS modulates spontaneous neural firing in the brain via sub-threshold alterations of membrane potentials [12,13]. Thus, tDCS is a purely neuro-modulatory intervention [13].

In previous tDCS studies, a 0.5–2 mA tDCS stimulation is often applied directly to the scalp via two 20-35 cm^2^ electrode pads (one anode and one cathode). The anode is placed on the scalp above the target region and the cathode is often placed on the scalp above the contralateral region, neck, or another area [11–14]. This stimulation method affects a large brain area between the anode and cathode [14,15]. In some cases, the target region may not even be included in the actual stimulated region depending on the locations of the electrodes. To overcome the drawbacks of this kind of large electrode pads, numerous new tDCS electrode montages have been developed to improve the guidance of the stimulation to the target region. These include high-definition tDCS [16], and dense electrode array tDCS [17–21]. In high-definition tDCS, the stimulation precision is improved by using different types of electrodes or electrode placement. Saturnino et.al. investigated the effects of several types of electrodes and electrode placement on the brain stimulation results, and recommended that using a 4 × 1 ring electrodes placement or two focal center surround ring electrodes can achieve better focality of stimulation [16]. In dense electrode array tDCS, *N* electrodes or *N* pairs of electrodes are positioned on the surface of the whole scalp according to a specific principle (e.g., Electroencephalogram (EEG) 10/20 or 10/10 system), and the stimulated target is located by using different current distributions on these electrodes (i.e., giving each electrode a specific current). Dmochowski and Guler pointed that when the current of each electrode is calculated using optimization algorithms under certain safety constraints, the stimulation via dense electrode array tDCS can be generally limited to the specific target region [17,18].

When using optimization algorithms, the selection of an optimization scheme (i.e., cost function) is a key problem. Different optimization schemes will induce different optimization results because of the varied optimization emphases in each scheme. Most dense electrode array tDCS simulation studies have used the minimum least squares (MLS) scheme to optimize the current distribution in each electrode, which minimizes the sum of squared error between estimated stimulation results and the expected one [17,20]. The MLS scheme must specify the strength and direction of the expected stimulation. The direction is often set as the normal or tangential direction of the location of stimulation. But the strength needs grid research, which sometimes can be very time consuming. Guler proposed a new optimization scheme involving the maximum electrical field strength (ME) inside the target region [18]. In comparison with the MLS scheme, the ME scheme can induce the maximum stimulation strength in the target region and only needs to specify the direction of the expected stimulation. However, these studies only examined optimization performance in the case of a single target stimulation. The performance of the MLS and ME schemes in multi-target stimulation scenarios is unknown. As individual brain functions often involve many brain regions, simultaneously stimulating two or more brain regions may have more impact on a specific brain function. Dmochowski proposed a MLS based optimization scheme that uses the EEG recording to give neural sources which generate this EEG recording a stimulation [20]. Although, this method can give neural sources a relatively precise stimulation without given any additional parameter such as the strength and direction of the expected stimulation, it can only apply to the situation that neural sources which generate the recorded EEG need to be stimulated. Ruffini and his colleagues first proposed the concept of multifocal transcranial current stimulation [21]. In this concept, all brain voxels are stimulated under a specific stimulation plan according to a given pattern from neuroimaging studies, such as a T-map in functional magnetic resonance imaging or a apparent diffusion coefficient map in diffusion tensor imaging [21]. However, this study only proposed a framework. The selection of an optimization scheme, specific parameter settings of each optimization scheme, and their effects on the optimization results were not given in detail.

Thus, in this study, we systematically investigated the optimization performance of the MLS and ME optimization schemes when applied to brain stimulation via a 64-electrode dense array tDCS system, described previously [17]. We examined the performance of the two schemes in three conditions: 1) single target stimulation optimization, 2) multi-target stimulation optimization, and 3) stimulation optimization under an activation by a specific task.

## MATERIALS AND METHODS

### Problem Description

Consider the five heterogeneous layers in the head model shown in Fig 1 in Supplementary Material. Each layer has a scalar conductivity field *σ*. Assuming that each layer is isotropic, for a steady current injected from the scalp to the brain (e.g. tDCS), the current density J of the brain tissue should have zero divergence. Thus, when applying the current through electrodes onto the scalp, the potential distribution *V* of the brain can be calculated by solving Laplace’s equation under a boundary value condition [22]:

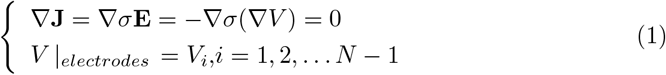

**Fig 1.**
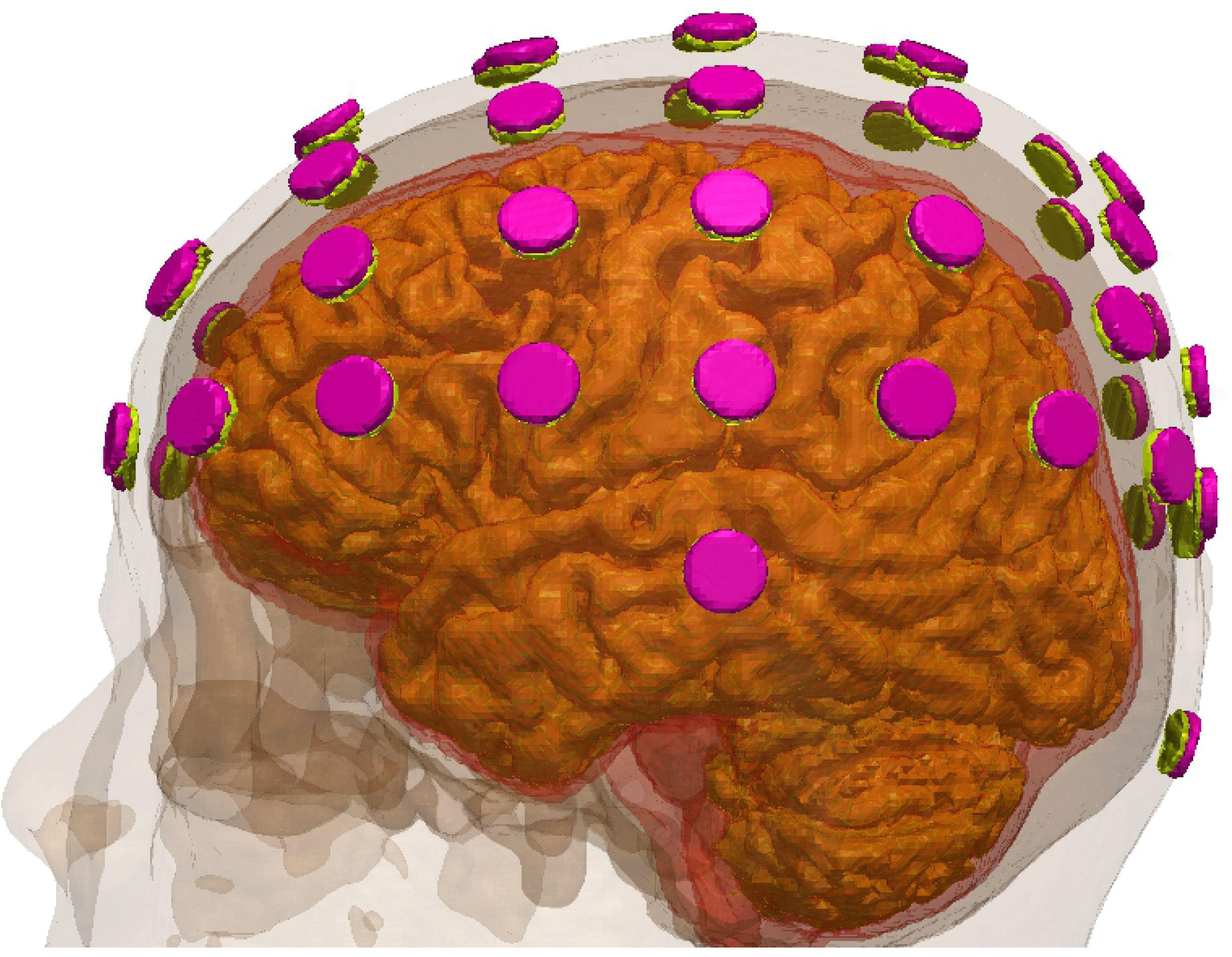
The constructed head model with five types of tissue, electrodes (pink), and conductive gel units (yellow).

Where **E** represents the electrical field distribution of the brain, *V_i_* represents the stimulation potential of the *i*th electrode area, and *N* is the number of electrodes. Eq 1 indicates that, given each a potential *V* for each electrode, the potential anywhere in the brain can be calculated by solving Laplace’s equation.

However, in general, the analytic solution of Eq 1 does not have a closed-form solution. Thus, in technical applications, analytic solutions are often replaced by numerical approximations. In this study, we used the finite elements method (FEM) [23] to calculate this numerical approximation. The FEM is the most widely used method for approaching this type of calculation. It firstly segments the whole brain volume into finite (usually a million or more) mutually disjointed elements (that usually form a tetrahedron). By solving the equivalent variational problem of Laplace’s equation for each element, FEM translates Eq 1 into a linear homogeneous equation system [23]:

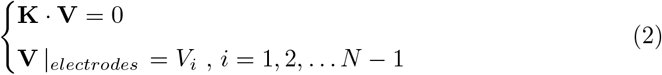

Where **V** = [*v*_1_, *v*_2_, …, *v_M_*]^T^ represents the potential of the nodes that are the vertices of these elements, *M* is the total number of nodes, and **K** is the coefficient matrix of the linear homogeneous equation system. The **K** is derived from the space coordinates and corresponding conductivity of each node. Eq 2 not only simplifies the problem described by Eq 1, but also markedly improves the speed at which solutions can be computed, making FEM a popular simulation method in many fields. Solving the potential of each node using Eq 2 enables the calculation of **V**, the corresponding electrical field **E**, and the current density **J** of each node in brain [22].

Consider that *N* electrodes are applied onto the surface of scalp layer of a head model. Assume that one of these electrodes is a cathode (reference electrode) and the other *N*-1 electrodes are anodes (freedom electrode). Application of a unit of stimulation (such as 1A or 1mA) at the ith anode when the other anodes are set to zero will produce a unit electrical field distribution matrix **a_*i*_**(**r**) throughout the brain volume with the space coordinates of each node, **r**, as a variable using FEM, which is only associated with the *i*th anode. According to the principle of linear superposition, if stimulation with an arbitrary current magnitude is applied to each anode, the electrical field distribution can be described as follows [17]:

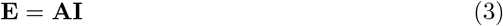

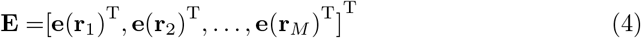

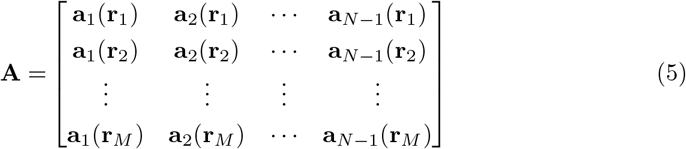

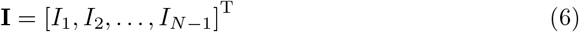

Eq 3 to Eq 6 imply that by calculating the distribution coefficient matrix **A** using FEM, the electrical field distribution **E** can be calculated as a straightforward linear solution. Thus, given a specific electrical field distribution **E**, we can use an optimization scheme to calculate a set of stimulation currents **I** that can generate an 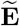 that will maximally approximate **E**.

### Optimization schemes

In this study, we investigated the performance of tDCS in a multi-object stimulation scenario using two optimization schemes: MLS and ME scheme.

#### Minimum Least Squares Scheme

The MLS optimization scheme is widely used in many research fields. Given a specific electrical field distribution **E**, the MLS optimization scheme is defined as follows [17]:

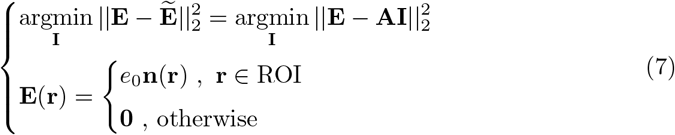

Where ∥ · ∥_2_ represents the 2-norm of the vector, *e*_0_ represents the expected electrical field strength inside the region of interest (ROI), and **n**(**r**) represents the expected normal direction vector at **r**. Eq 7 indicates that the goal of the MLS scheme is the minimization of the squared error between the estimated electrical field 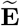 and the expected electrical field **E**. Considering the safety constraint that the absolute current value of each electrode should not exceed *I_max_* and the absolute current value flowing into the cathode should equal the sum of the current value flowing out of the anodes (i.e., |*I_cathode_*| = |∑*I_anodes_*|), the MLS optimization scheme in Eq 7 can be adjusted as follows [22] to minimize the squared error under some specified safety constraints:

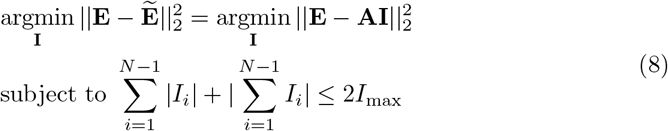

As decribed in [17], if current satisfies this sum form of current constrains, then |*I_i_*| ≤ *I*_max_, and |∑ *I_i_* ≤ *I*_max_

In FEM, the number of nodes *M* is usually very big (i.e., 10^6^-10^7^ or greater). Thus, direct optimization using Eq 8 can be both time and source consuming. Further simplification of Eq 8 is need. The least squares can be extended as:

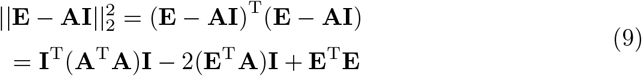

Where **A^T^A** is a (*N* – 1) × (*N* – 1) matrix, **E^T^A** is a 1 × (*N* – 1) vector (*N* ≪ *M*), and **E^T^E** is a constant. If we let **W**_*aa*_ = **A^T^A**, **w**_*ea*_ = **E^T^A**, and *w_ee_* = **E^T^E**, Eq 8 can be simplified as:

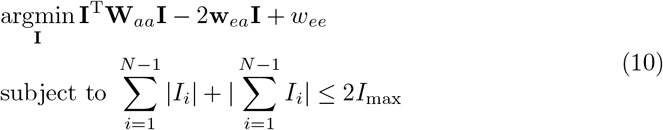

As **A** is already calculated using FEM, **E** is given, and **W**_*aa*_, **w**_*ea*_, and **w_ee_** can be pre-calculated, optimization using Eq 10 can be completed in seconds.

#### Maximum Electrical Field Strength inside the ROI

In study [18], Guler stated that a MLS scheme should specify the strength and direction of the expected electrical field **E**. In general, researchers tend to select the direction as the normal or tangential direction [17,18,20]. However, the strength of the expect electrical field *e*_0_ is very important, as it will decide not only the stimulation strength of 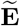, but also the form of 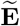 as well (Fig 6). Thus, Guler proposed a new optimization scheme, the maximum current density **J** inside the ROI, which only requires specification of the direction of the expected electrical field **E**. As **J** = *σ***E**, where *σ* is the conductivity of medium, the maximum current density inside the ROI is equivalent to the maximum electrical field inside the ROI, i.e., ME The math formulation of the ME scheme is defined as follows [18]:

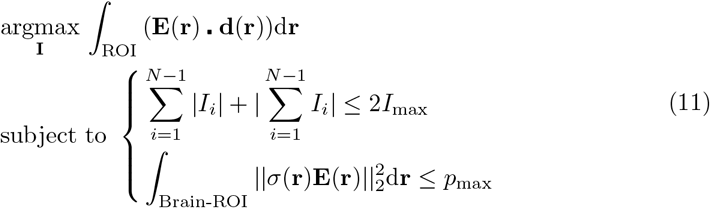

Where 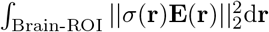 represents the total power outside the ROI that, with regards to safety considerations, should not be larger than *p*_max_. · represents the dot product operation. Eq 11 shows that the goal of the ME scheme is to maximize the sum of the electrical field in the target ROI under the safety constraint. According to Guler [18], Eq 11 can also be simplified. The ∫_ROI_ (**E**(**r**)**.d**(**r**))d**r** can be discretized as:

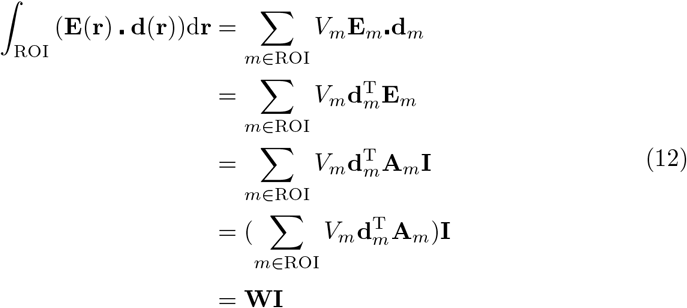

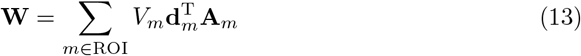

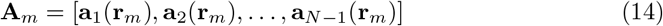

Where *V_m_* represents the volume of the *m*th elements. If we assume uniform distribution of the nodes that construct elements in the brain, then *V_m_* can be approximately treated as a constant. Thus, Eq 13 can be approximately represented as:

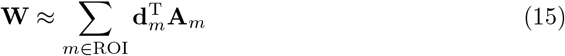

Similarly, 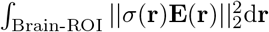 can be extended as:

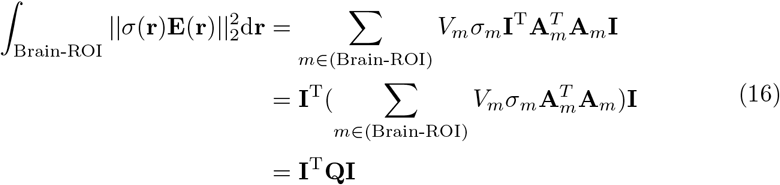

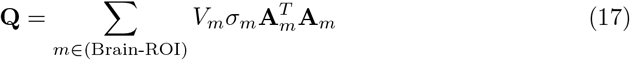

Eq 11 can be simplified as [18]:

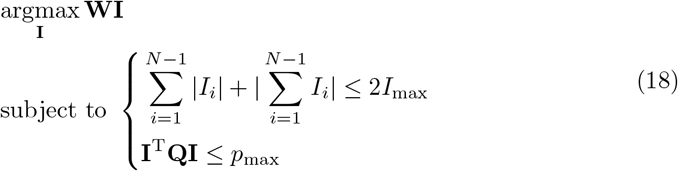

Also, **W** and **Q** can be pre-calculated. Optimization operations using Eq 18 can be also completed in seconds.

Eq 10 and Eq 11 give the basic formulation of MLS and ME optimization scheme in single target case. However, when we directly used these two schemes in multi-target (i.e. taking all target stimulation brain regions as a whole region), the simulation results were not be the one we desired (Fig 5 & 6). To correctly guide stimulations to target regions in the multi-targets case, we propose the weighted ME and weighted MLS optimization scheme in next two subsections.

### The Weighted ME Scheme

Using the ME scheme described in Eq 18, the **W** can be rewritten as,

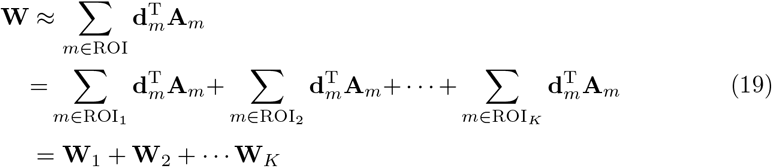

Thus, the basic weighted ME scheme can be defined as follows:

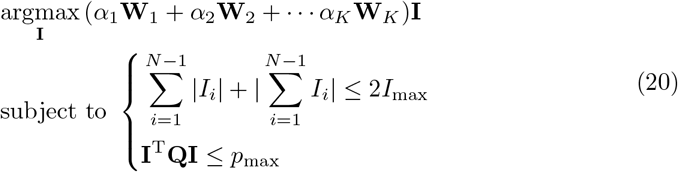

Where *α_j_* represents the weight assigned to the *j*th ROI and satisfies *α_j_* > 0, *j* = 1, 2,… *K*. To account for variations in the sizes of the ROIs, before applying the expected weights to each ROI, the original weight of each ROI, which is mainly proportional to the size of the ROI, should be equalized. Thus, the *α_j_* can be defined as follows:

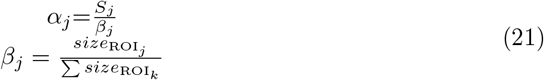

Where 1/*β_j_* is the equalization coefficient of the *j* th ROI and *S_j_* should be calculated according to specific requirements. Because the size of a ROI is not the only factor that affects the original weight, for accurate calculation, *β_j_* can be redefined as:

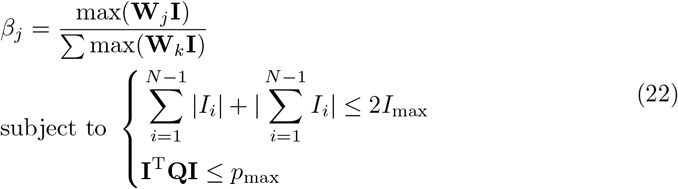

From Eq 22, for accurate calculation of *β*, single target optimization for each ROI using the ME scheme should be computed first. As this can be time consuming, *β* as defined in Eq 21 is sufficient for general use.

If the goal is to stimulate all ROIs with a similar strength, a strength distribution constraint should be added to Eq 20:

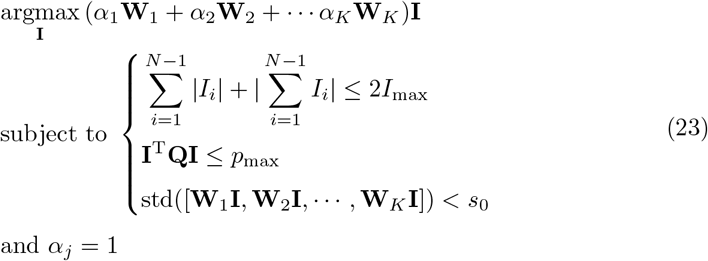

Where std(·) represents the standard deviation of a vector and *s*_0_ > 0 is the maximum standard deviation. The smaller the *s*_0_, the more similar the stimulation strength among the ROIs. The main effect of Eq ?? is to constrain the differences between ROI stimulation strengths to a relatively small range.

If primary electrodes that maintain the stimulation for each ROI need to be injected with a similar current, the weighted ME scheme should be rewritten as:

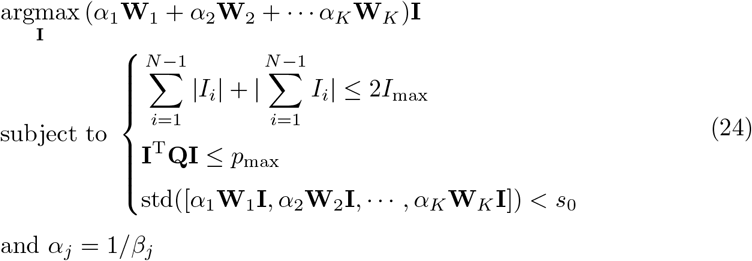

In Eq 24, *α* equals to the equalization coefficient. Weighting *α* for each ROI is equivalent to setting each ROI to a similar size. However, simply weighting the ROIs with *α* is not sufficient. In this situation, if we further constrain the differences among each ROI’s weighted stimulation strength to a relatively small range, the current injected into each primary electrode will be similar.

If the goal is to stimulate some ROIs with greater strength and other ROIs with less strength, an additional current distribution constraint should be added:

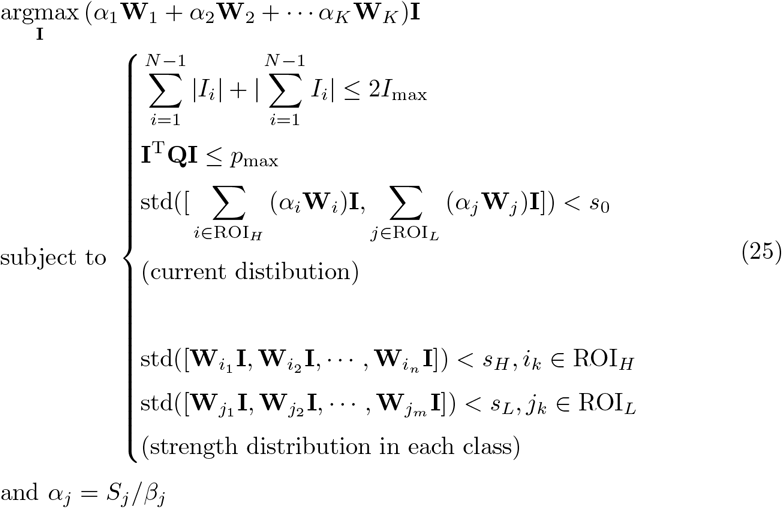

Where the *S_j_* needs to be calculated according to specific distribution requirements, and in each ROI class (e.g., high stimulation class and low stimulation class), *S_j_* should be same. The subscript H and L represent the High and Low, respectively. If a class has only one ROI, the strength distribution constraints for that class can be removed from Eq.25. The current distribution constraint added to Eq 25 is necessary because the original strength ratio is generally not equal to the ratio we desired (e.g., 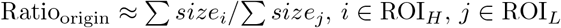). Eq 25 first distributes the injected current for each class according to *α*, which causes the strength ratio between each ROI class to approximate the desired ratio. After that, each ROI’s stimulation strength is distributed in each class according to the corresponding strength distribution constraint.

### The Weighted MLS Scheme

From the MLS scheme described in Eq 10, **E** can be rewritten as:

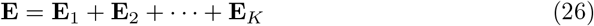

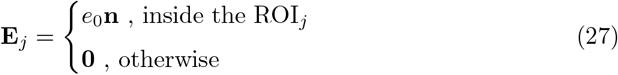

Thus, the weighted MLS scheme can be defined as follows:

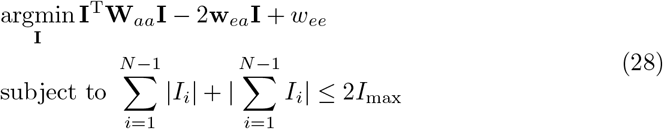

Where,

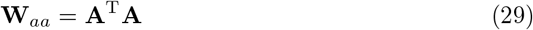

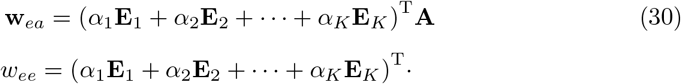

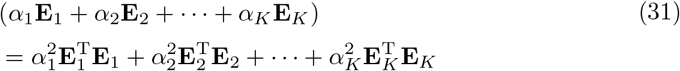

Unlike weighted ME schemes in which strength distribution and current distribution constraints are needed, only the weight *α* is needed in a weighted MLS scheme. The *α* needs to be carefully calculated according to the specific stimulation mode. Theoretically, the larger the *α* and the larger the size of a ROI, the larger the error minimization priority that a weighted MLS scheme will give to that ROI. If the goal is to obtain a weighted MLS scheme that will give equal priority to each ROI, equalization should be first conducted according to the size of each ROI. The equalization weight *β* has the same definition as in Eq 21. However, if there exists a ROI that is too small with respect to the other ROIs, the equalization weights for this ROI will be too large. This may cause an ‘overfitting’ problem where the optimization algorithm will only minimize this ROI and ignore the others. In this situation, it may be necessary to appropriately increase the weights of the other ROIs. Finally, the weighted MLS scheme is very strongly affected by the expected electrical field strength *e*_0_. Thus, it is important to search for an appropriate *e*_0_, but this process can be time consuming.

### Experimental Setup

#### Realistic Head Model

We used high resolution T1-weighted MR brain images of a healthy female participant from our previous study [24] to construct a realistic stimulation head model. These MR images were acquired using a 3T Philip scanner at the Department of Radiology, Xijing Hospital, The Fourth Military Medical University (voxel size: 1*mm* × 1*mm* × 1*mm*; resolution: 256 × 256 × 192). We used FSL software (https://fsl.fmrib.ox.ac.uk/fsl/) to segment brain tissues into scalp, skull, cerebrospinal fluid (CSF), gray matter (GM), and white matter (WM). Briefly, brain tissues were first segmented into scalp, skull, and brain areas using the brain extraction tool (BET) function. Then, the brain area section was further segmented into CSF, GM, and WM using the FMRIB’s automated segmentation tool (FAST) function.

After segmentation, the five tissues masks were imported into simpleware software (https://www.synopsys.com/simpleware.html) to generate the final head model (Fig 1). We performed manual correction of the tissue masks using the ScanIP module. Then, we used the ScanCAD module to place 64 cylinders with a diameter of 12 mm and depth of 2 mm [17], modeled as electrodes, on the scalp according to the international EEG 10/10 system. Additionally, we placed 64 cylinders with a diameter of 12 mm and depth of 1 mm, modeled as conducting gel [17], on the scalp with the upper and lower surface directly contacting each electrode and scalp surface, respectively. Finally, we translated the entire head volume consisting of 133 entities (64 electrodes + 64 conduction gel units + 5 types of brain tissue) into a finite element mesh using the ScanFE module. The conductivity of each entity and the other parameters required in this study are detailed in Table 1.

**Table 1.** Parameters list of head model

| Name | Value | Note |
| --- | --- | --- |
| Scalp | 0.333 S/m | [25] |
| Skull | 0.0083 S/m | [25] |
| CSF | 1.79 S/m | [25] |
| GM | 0.333 S/m | [25] |
| WM | 0.333 S/m | [25] |
| Electrode | 5.9e7 S/m | [17] |
| Conductive Gel | 0.3 S/m | [17] |
| $I_{max}$ | 2 mA | [17] |
| $p_{max}$ | 1e-7 A<sup>2</sup>/m (about 1e-6 V<sup>2</sup>·m) | [18] |

#### Performance Metrics

To evaluate the performance of each optimization scheme, we considered two aspects: intensity and focality. To assess the stimulation intensity of a given ROI, we used the total electrical field strength inside the ROI, defined as the cost function **WI** of the ME scheme. To evaluate the focality of the stimulated ROI, we used the cross correlation coefficient (CC) defined in Ruffini’s study [21]:

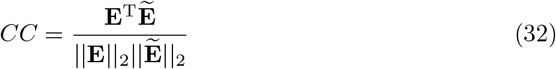

#### Simulation Implementation

In this study, we used COMSOL software (https://www.comsol.com/) to solve Laplace’s equation using FEM. We used MATLAB (https://www.mathworks.com/) to implement the optimization scheme. We selected the Cz point in the EEG 10/10 system to be the cathode (reference electrode). Because of the generally vere long time of tDCS stimuli, we predicted steady-state fields without much concern for the charging/discharging effect of tissue capacitance (i.e., using quasistatic solution). The procedure is shown in Fig 2.

**Fig 2.**
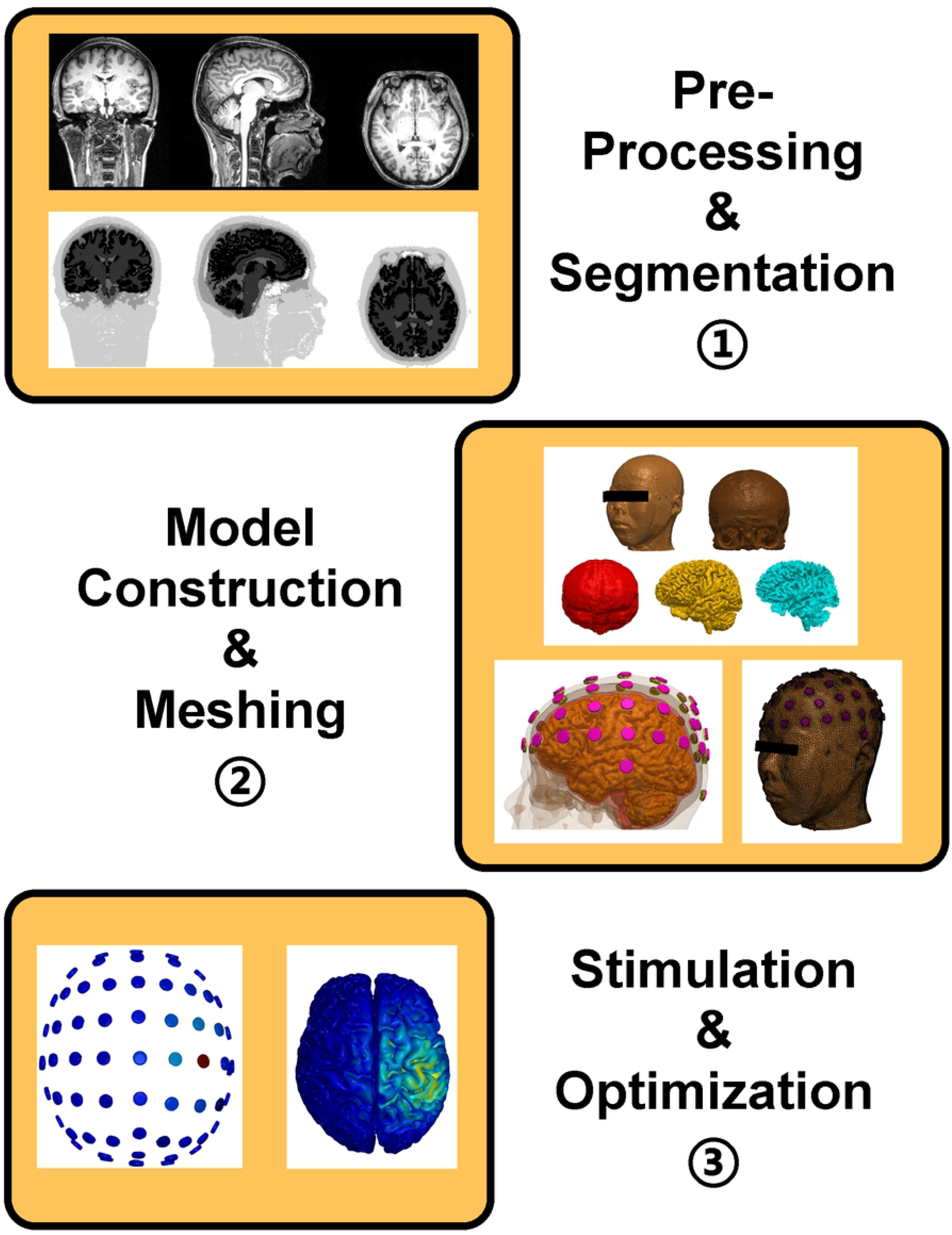
The study procedure. 1) Subject T1-weighted MR images were first segmented into five brain tissue masks. 2) We constructed a head model by meshing the five tissue masks. 3) FEM stimulation and current optimization was performed for each electrode using the meshed head model.

## RESULTS

### Single Target Case

Fig 3 shows the single target optimization results of two selected ROIs based on an anatomical automatic labeling (AAL) atlas [26] for two optimization schemes (left column for MLS and right column for ME). The two ROIs were, from top to bottom, the right precentral gyrus (R PreCG) and anterior cingutate cortex (ACC), respectively. The ACC locates deeper in brain than PreCG. As shown in Fig 3, for a single target case, the MLS scheme had higher focality (i.e., more concentrated energy distribution in stimulation region) but lower intensity (i.e., lighter color in stimulation region) compared with the ME scheme. The detailed performance characteristics are given in Table 2. From Table 2, we can see that, when the depth of the stimulated region increased (PreCG to ACC), the intensity decreased for both MLS (0.0529 vs 0.0478) and ME schemes (0.0618 vs 0.0571). For the MLS scheme, the expected electrical field strengths were set to 2.0 V/m and 1.6 V/m for the PreCG and ACC, respectively, as shown in Fig 4D

**Fig 3.**
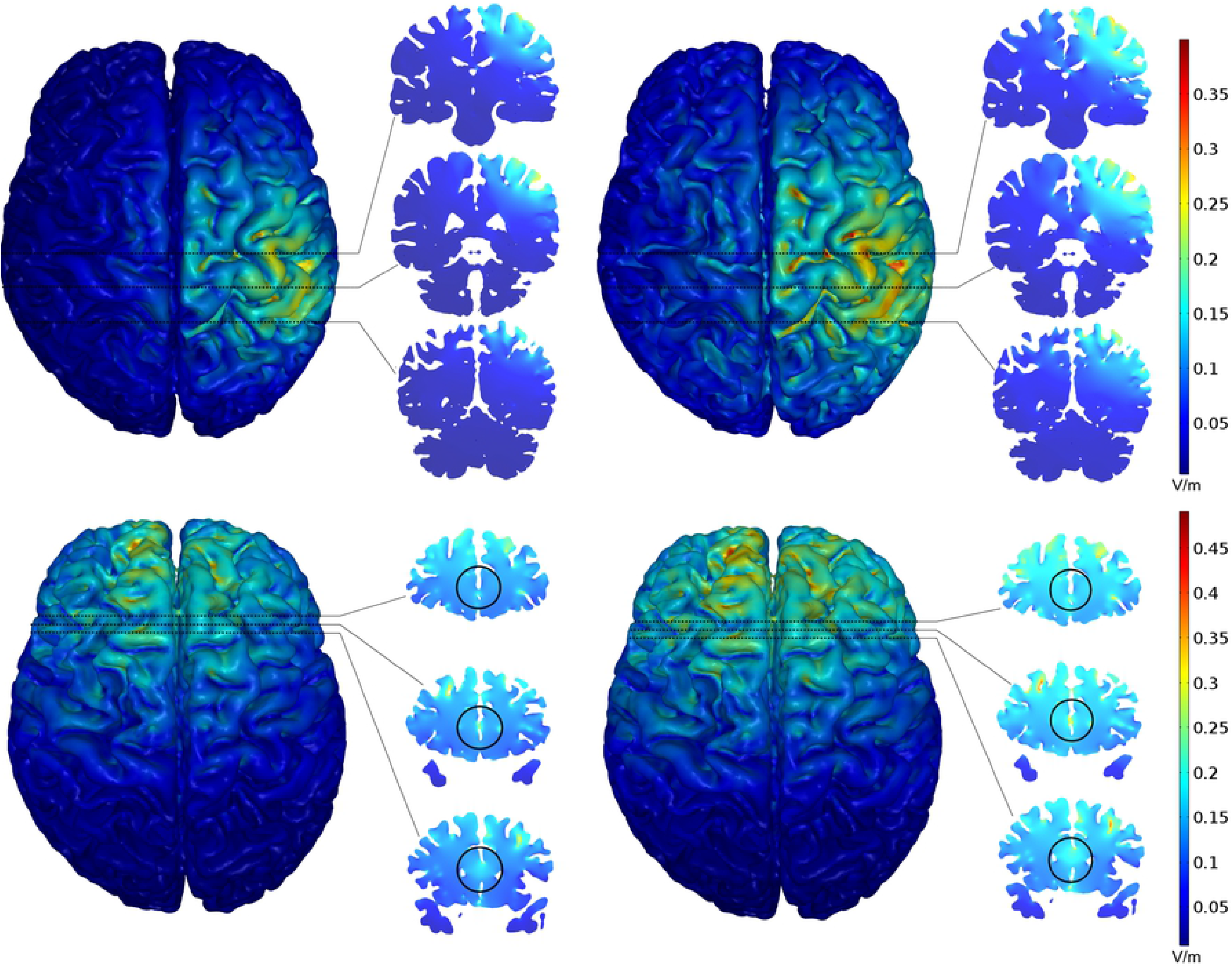
The optimization results of the MLS scheme (left) and ME scheme (right) for R PreCG (top) and ACC (bottom).

**Table 2.**
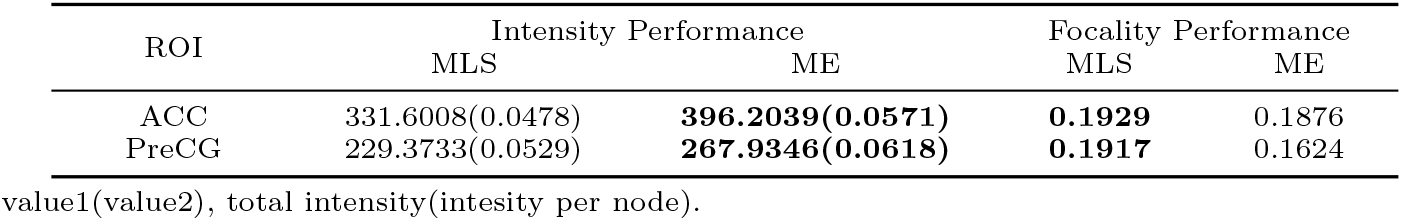
The performance of MLS and ME scheme

| ROI | Intensity Performance |  | Focality Performance |  |
| --- | --- | --- | --- | --- |
|  | MLS | ME | MLS | ME |
| ACC | 331.6008(0.0478) | **396.2039(0.0571)** | **0.1929** | 0.1876 |
| PreCG | 229.3733(0.0529) | **267.9346(0.0618)** | **0.1917** | 0.1624 |

value1(value2), total intensity(intensity per node).

**Fig 4.**
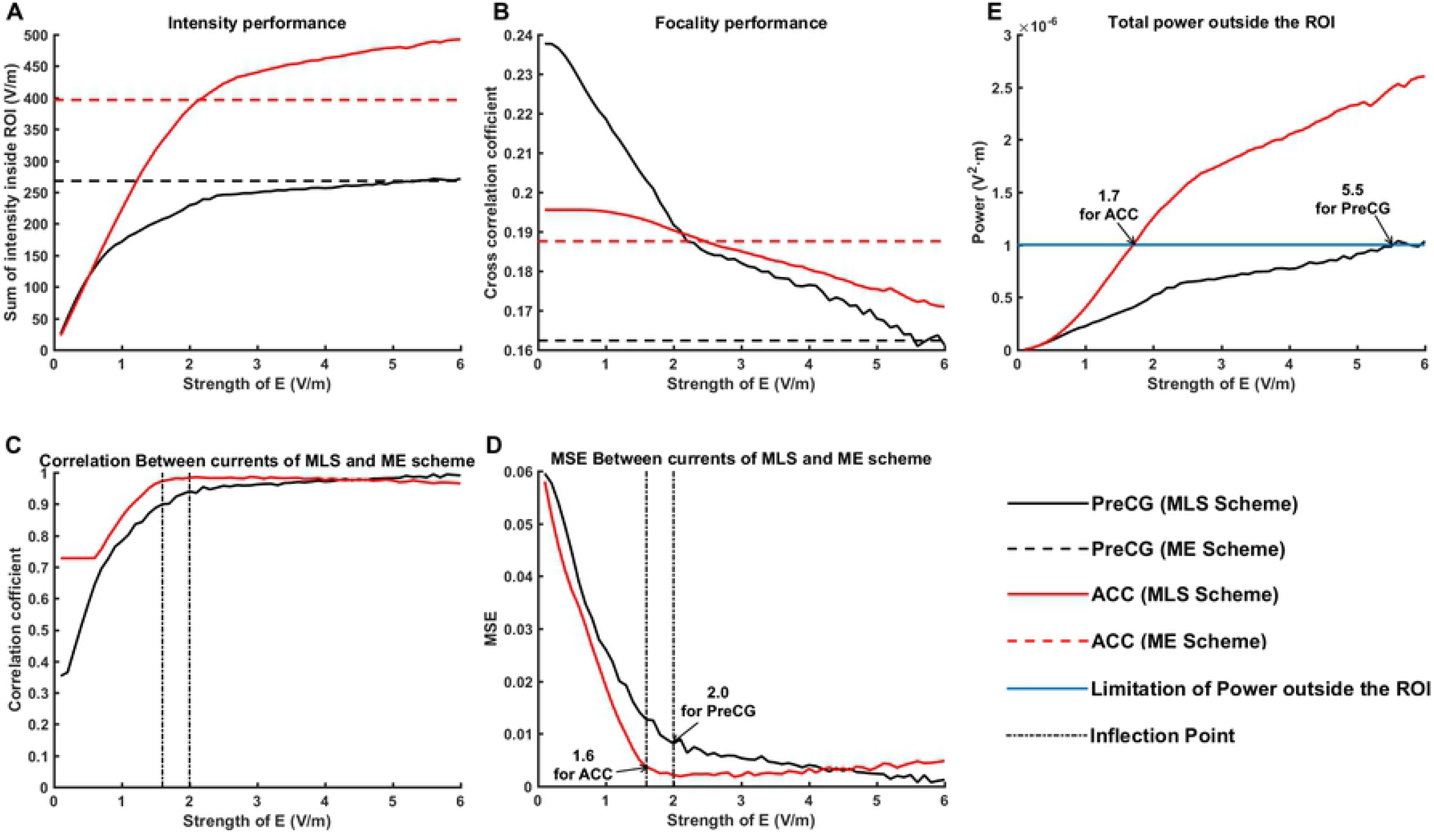
Features of the MLS scheme changed according to the expected electrical field strength. A) Intensity performance; B) Focality performance; C) The correlation coefficient between currents in the MLS and ME schemes; D) The MSE between currents in the MLS and ME schemes; E) The total power outside the ROI.

As displayed in Fig 4, when we gradually increased the strength of the expected electrical field in the MLS scheme, we found that 1) the intensity performance increased and the focality performance gradually decreased, which demonstrated a trade-off relationship between focality and intensity (Fig 4A & 4B). 2) The focality and intensity values gradually approximated those in the ME scheme when power constraints existed outside the ROI (*P*_Brain-ROI_ ≤ 1e-6 V^2^·m). For the ROI on the grey matter on the surface of the brain (e.g., PreCG), the performance of the MLS scheme was nearly equal that of the ME scheme when the strength of the expected electrical field was set at the maximum value (5.5 V/m, Fig 4E). The maximum value represents the maximum strength to which the expected electrical field *e*_0_ can be set. Values larger than this will cause optimization results which do not satisfy the power constraint (i.e., *P*_Brain-ROI_ > *p*_max_). For deeper ROIs in the brain (e.g., ACC), the MLS scheme had better focality but lower intensity compared with the ME scheme when the strength of the expected electrical field was set at the maximum value (1.7 V/m, Fig 4E). 3) When the expected electrical field strength gradually increased to the maximum allowed value, Person’s correlation coefficient between the currents calculated by MLS and ME schemes gradually increased to one (Fig 4C). These findings indicate that the performance of the MLS scheme might be limited to that of the ME scheme when the strength of the expected electrical field is set at the maximum value.

### Multi-Target Case

The first column in Fig 5 illustrates the multi-target optimization results for three selected ROIs based on the AAL atlas for ME optimization schemes. The three ROIs were the left superior part of the frontal gyrus (L SFG), R PreCG, and left middle part of the occipital gyrus (L MOG). From the figure we can see that, when using ME scheme (i.e., *α*_SFG_ = *α*_PreCG_ = *α*_MOG_=1), the L SFG received the greatest stimulation, the R PreCG received medium stimulation, and the L MOG received almost no stimulation.

**Fig 5.**
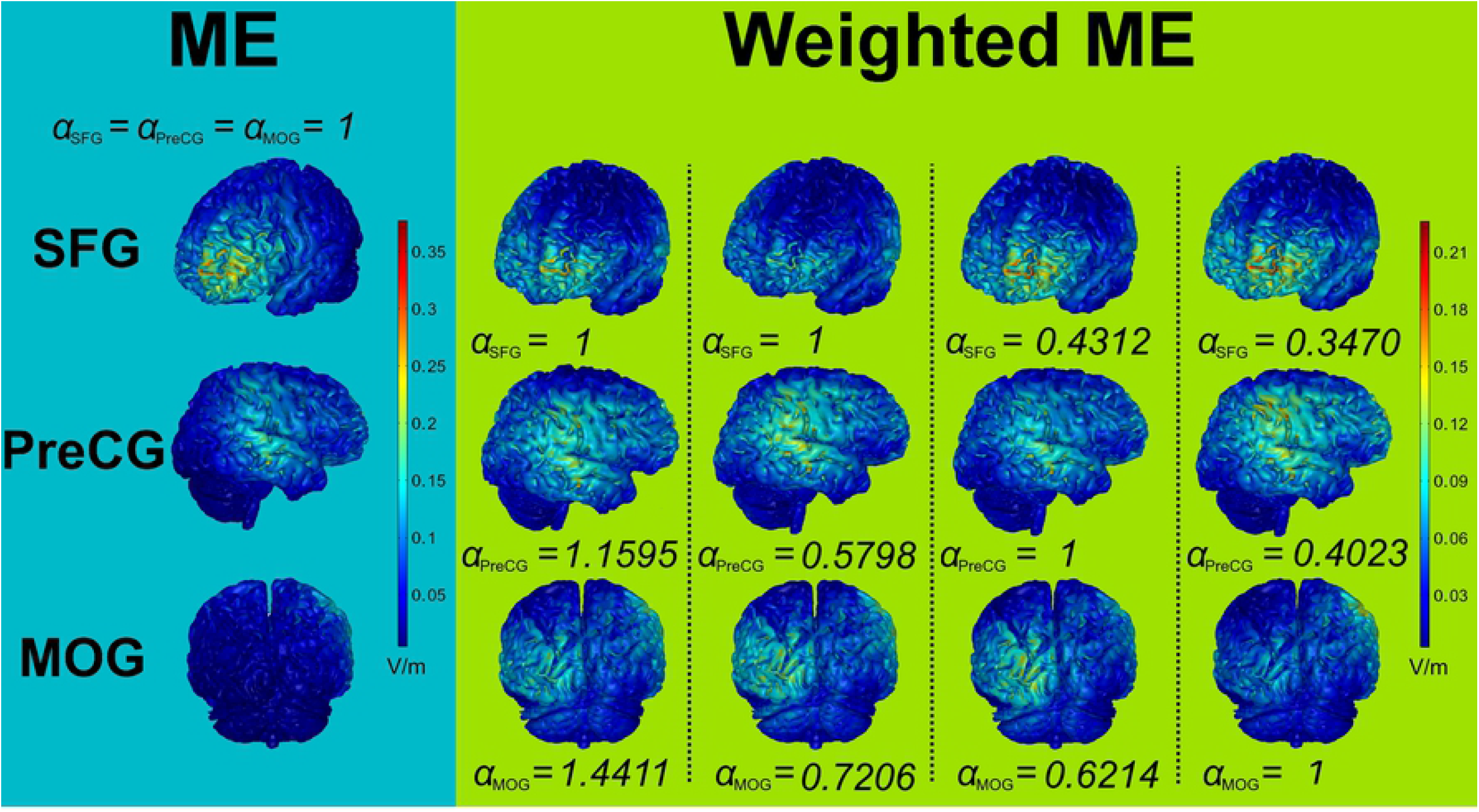
The optimization results of ME scheme (the first column) and results of weighted ME scheme for four stimulation modes: 1) all three ROIs have similar stimulation strengths (the second column), and 2) L SFG (the third column), 3) R PreCG (the fourth column), and 4) L MOG (the fifth column) have low stimulation strengths, respectively, while the other two ROIs have high stimulation strengths.

Table 3 displays the single target intensity performance of the ME scheme for these three ROIs. The results showed that under the same constraints, the L SFG had the largest intensity performance while the L MOG had the smallest intensity performance, mainly because it had the lowest number of nodes inside the ROI (i.e., smaller ROI). Similar to the ‘overfitting’ problem in machine learning field [27], the optimization algorithm might tend to give the L SFG a larger stimulation strength to achieve the goal of maximizing the sum of the overall electrical field strength across the three ROIs. Thus, in cases where different stimulation strengths are desired for different ROIs (i.e., stimulating in a different given mode), a weighted ME scheme is needed. As shown from the second column to the fifth column in Fig 5, when giving each ROI an appropriate weighting *α*, the weighted ME scheme can be useful for applying many stimulation modes. For example, when setting *α* = 1/*β* and using Eq.24, all of three ROIs would have similar intensity level of stimulation (the second column in Fig 5). On this basis, when multiplying *α* by [0.6, 0.3, 0.3], [0.3, 0.6, 0.3] and [0.3, 0.3, 0.6] respectively, the ROI_SFG_, ROI_PreCG_ and ROI_MOG_ would have lower intensity level of stimulation respectively, and the other two ROIs would have higher intensity level of stimulation (the third column to the fifth column in Fig.4). The *α* in Fig 5 were normalized. The derivation of *β* can be found in Eq 21 or Eq 22..

**Table 3.** The intensity performance of ME scheme on selected ROIs (Single Target)

| ROI | Number of nodes<br>in head model | Intensity |
| --- | --- | --- |
| L SFG | 4458 | 276.0214 |
| R PreCG | 4337 | 267.9346 |
| L MOG | 3278 | 224.7673 |

Fig 6 illustrates the multi-target optimization results for three selected ROIs based on AAL atlas for MLS optimization schemes. From the figure we can see that, when using MLS scheme and setting *e*_0_ ∈ [0.5, 2.0]V/m, all of three ROIs would have similar intensity level of stimulation and the total intensity level would increase gradually with the *e*_0_ increased. When setting *e*_0_ ∈ (0.5,3.7]V/m, the intensity level of ROI_MOG_ would decrease gradually with the *e*_0_ increased. Thus, changing *e*_0_ would not achieve the precise stimulation energy distribution. For other stimulation mode, a weighted MLS scheme is also needed. As shown from the second column to the fifth column in Fig 7, when setting *α* = 1/*β* and using Eq 28 to Eq 31, all of three ROIs would have similar intensity level of stimulation (the second column in Fig.6). On this basis, when multiplying *α* by [0.3, 0.6, 0.6], [0.6, 0.3, 0.6] and [0.6, 0.6, 0.3] respectively, the ROI_SFG_, ROI_PreCG_ and ROI_MOG_ would have lower intensity level of stimulation respectively, and the other two ROIs would have higher intensity level of stimulation (the third column to the fifth column in Fig 7). Also, the *α* in Fig.6 were normalized.

**Fig 6.**
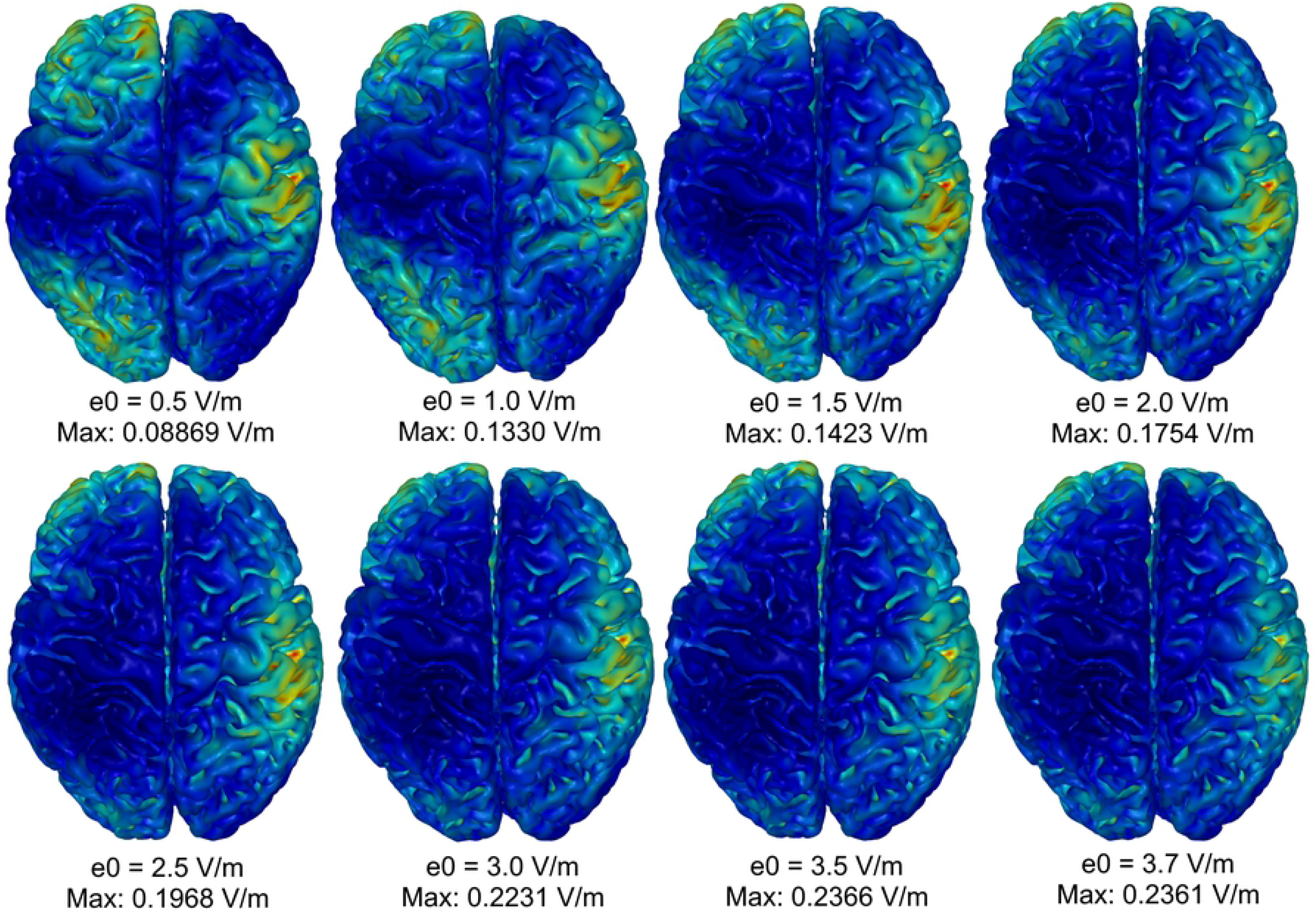
The optimization results for the MLS scheme changed with the expected electrical field strength *e*_0_. When *e*_0_ < 3.7 V/m, the total power outside the ROI was larger than 10^−6^ V^2^·m.

**Fig 7.**
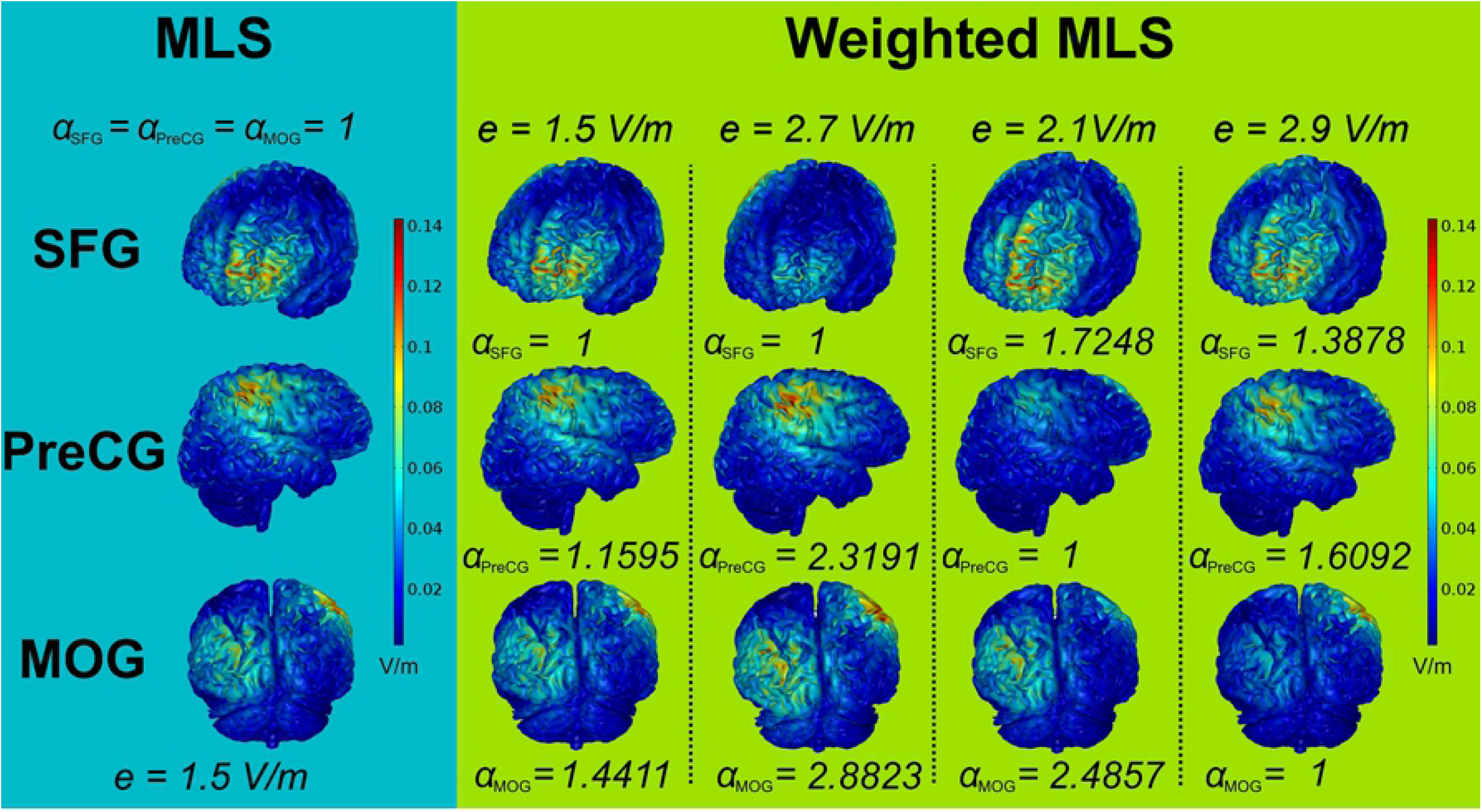
The optimization results of MLS scheme (the first column) and results of weighted MLS scheme for four stimulation modes: 1) all three ROIs have similar stimulation strengths (the second column), and 2) L SFG (the third column), 3) R PreCG (the fourth column), and 4) L MOG (the fifth column) have low stimulation strengths, respectively, while the other two ROIs have high stimulation strengths.

Compared Fig 5 with Fig 7, the energy distribution of weighted MLS scheme in each ROI was more concentrated than the one in weighted ME scheme, but the total intensity level was lower than the weighted ME scheme (colorbar in figures). Thus, similar with single target case, the weighted MLS scheme had better focality but a lower intensity compared with the weighted ME scheme.

### Multi-Target Under a Specific Task

We used activation T-map generated from functional MRI data during a specific space working memory task (SWMT) in 36 healthy humans as a reference for generating a stimulation mode according to the activation strength. The SWMT activation T-map is displayed in Fig 8 and the detailed activation ROI information is given in Table 4.

**Fig 8.**
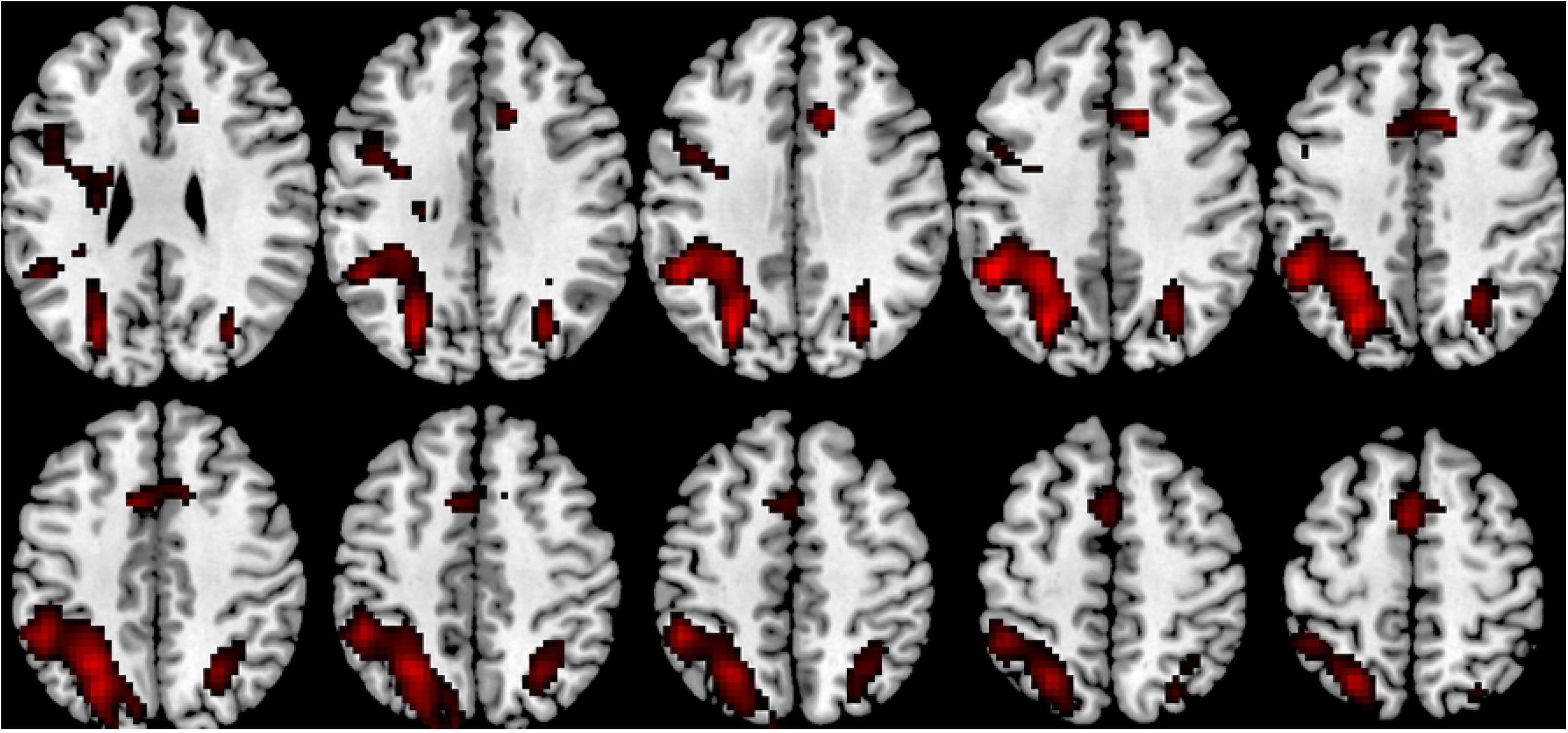
The SWMT activation T-map.

**Table 4.** The cluster information of SWMT activation T-map

|  | Region | Number of<br>nodes<br>in head models | Peak T<br>value |
| --- | --- | --- | --- |
| ROI 1 | R OG, R AG | 1251 | 7.3 |
| ROI 2 | L PreCG, L IFG | 429 | 5.7 |
| ROI 3 | L IPG, L SPG,<br>L OG | 5283 | 8.5 |
| ROI 4 | SMA, ACC | 1342 | 7.4 |

OG = occipital gyrus, AG = angular gyrus, PreCG = precentral gyrus, IFG = inferior frontal gyrus, IPG = inferior parietal gyrus, SPG = superior parietal gyrus, SMG = supplemental motor area and ACC = anterior cingulate cortex. L = left and R = right.

We optimized two stimulation modes using weighted ME and weighted MLS scheme, respectively. For the weighted ME scheme, we used a regional stimulation mode: each ROI had a stimulation strength sum that was proportional to the corresponding peak T value. The detailed parameters for the weighted ME scheme are shown in Fig 9 and the results are shown in Fig 9 and Table 5. The derivation of parameters used in the weighted ME optimization scheme can be found in Appendix. The results shown that the calculated parameters shown in Fig 9 were effective in evenly distributing the electrical field stimulation between each ROI (Table 5), indicating the correctness of our proposed method.

**Fig 9.**
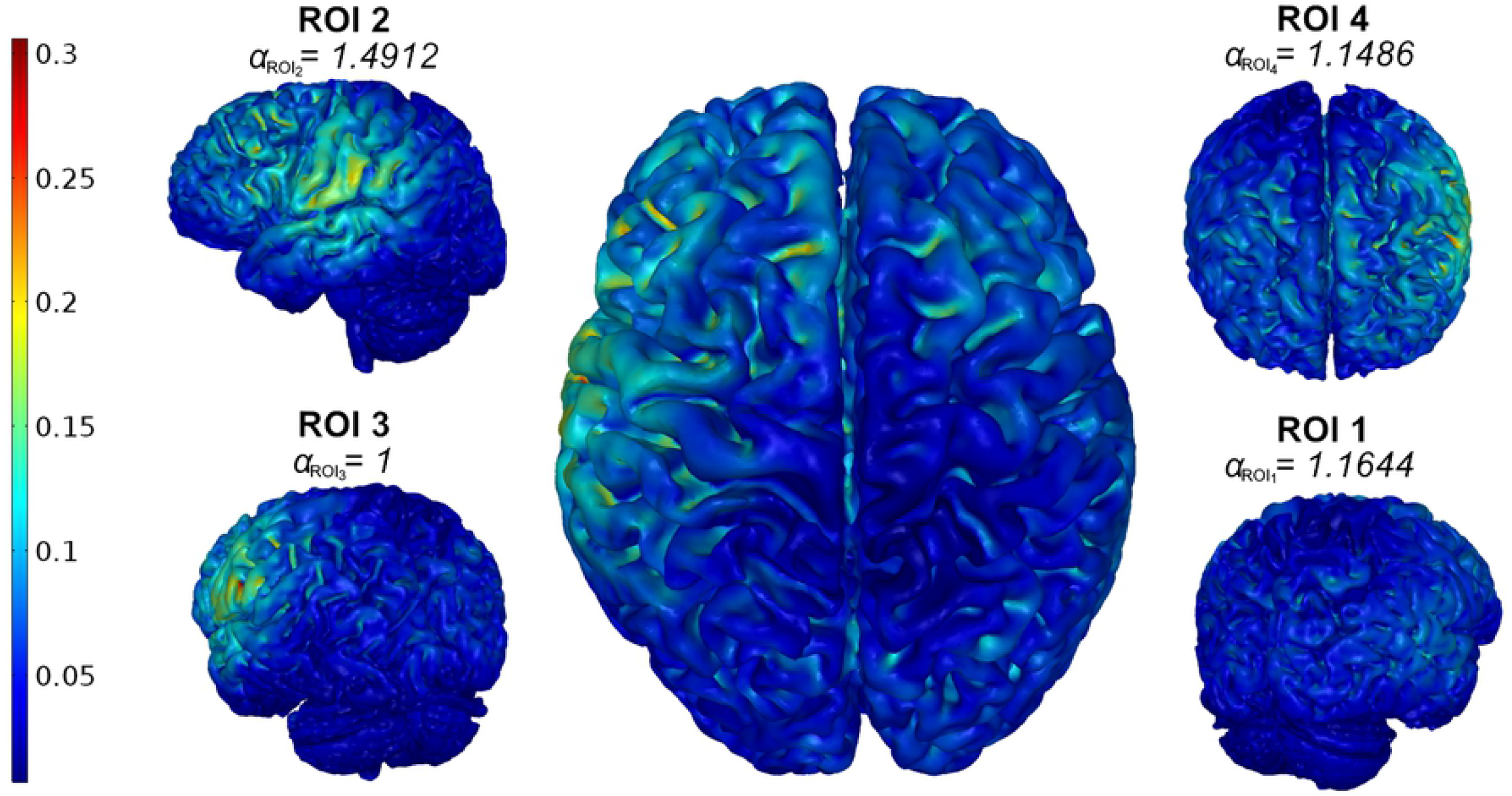
The optimization results of a regional stimulation mode using weighted ME optimization scheme.

**Table 5.** The optimization results of the weighted ME scheme

|  | ROI 1 | ROI 2 | ROI 3 | ROI 4 |
| --- | --- | --- | --- | --- |
| Peak T value | 7.3 | 5.7 | 8.5 | 7.4 |
| Total stimulation strength (V/m) | 26.9126 | 20.9440 | 31.4557 | 27.2971 |
| Ratio | **7.300** | **5.6810** | **8.5323** | **7.4043** |

The second stimulation mode directly used T values in the SWMT T-map as the weight of each voxel’s expected stimulation strength. We kept T values larger than 4.5 (*p* < 0.05, familywise error corrected). The others were set to zero and then normalized using the maximum T value in the map. This stimulation mode was not suitable for the weighted ME scheme, which was regional based. Before optimization, we performed equalization according to the size of each ROI (e.g., number of nodes in the head model). The detailed parameters are shown in Table 6. The result of the optimization is shown in Fig 10 and Table 7 with *e*_0_ selected as 1.5 V/m.

**Table 6.**
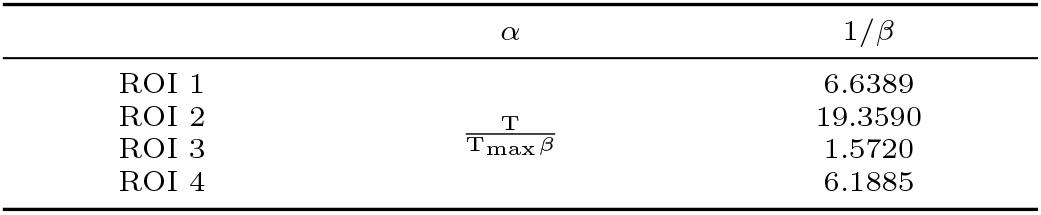
Parameters setting for the weighted MLS schemes

**Fig 10.**
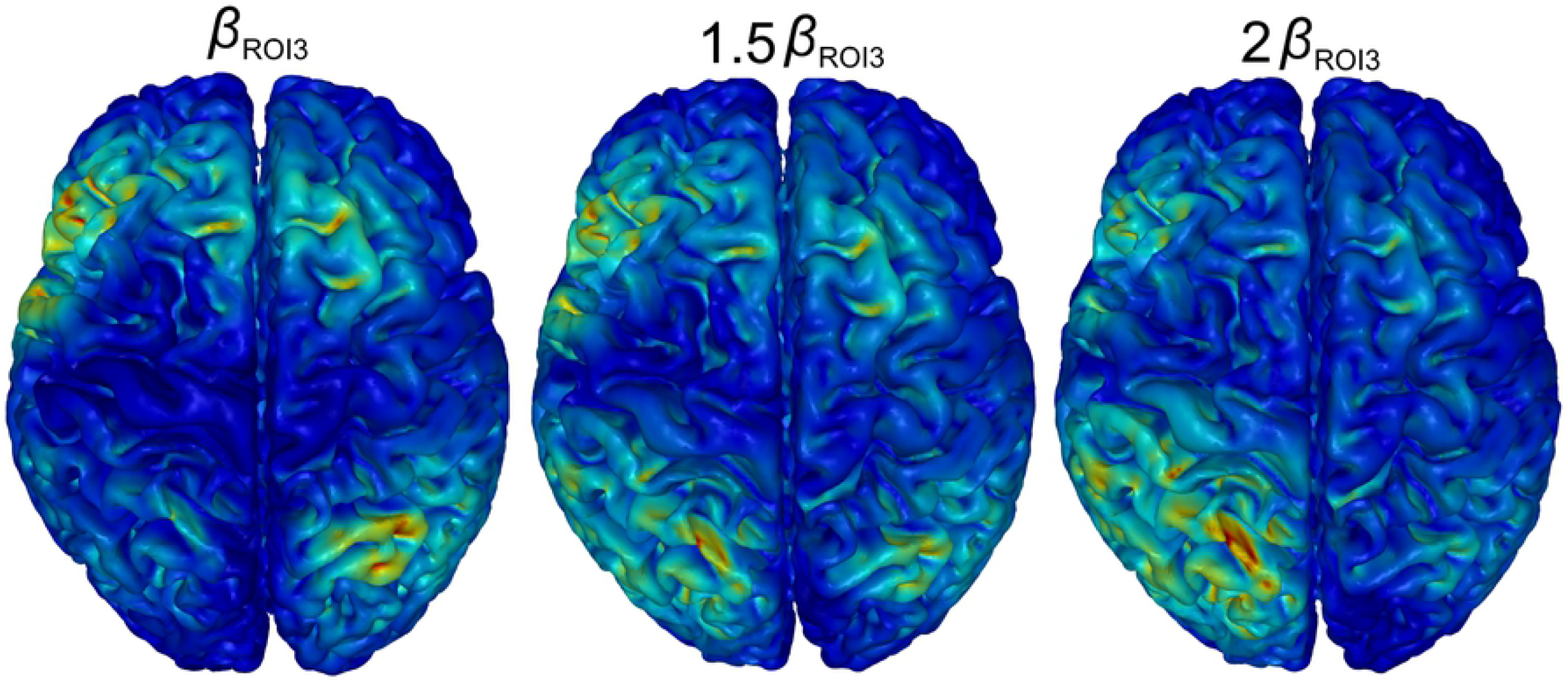
The optimization results of a voxel-wised stimulation mode using weighted MLS optimization scheme.

**Table 7.** Cross correlation coefficient of weighted MLS schemes

| | $\beta_{ROI3}$ | $1.5\beta_{ROI3}$ | $2\beta_{ROI3}$ |
| --- | --- | --- | --- |
| ROI 1 | 0.2028 | 0.2127 | **0.2087** |
| ROI 2 | 0.0715 | 0.1130 | **0.1450** |
| ROI 3 | 0.1409 | 0.1898 | **0.2197** |
| ROI 4 | 0.1598 | 0.1839 | **0.1878** |
| Outside ROIs | 0.1279 | 0.1587 | **0.1762** |

## DISCUSSION

We systematically investigated the optimization performance of two optimization schemes, MLS and ME, using a 64 electrode dense array tDCS system. We first investigated the optimization performance of two optimization schemes in the case of single target, to verify and complement the results of previous studies. Second, we investigated the optimization performance of two optimization schemes in the case of multi-target, to expand these two schemes to multi-target optimization. Finally, we used stimulation of a SWMT activation T-map to further validate our proposed expanding methods for the two optimization schemes. We discuss the corresponding results in detail below.

### Single Target Case

The MLS scheme had been widely used in many brain stimulation studies [17,20]. As its definition can be decomposed into a simple form Eq 10, the processing time can be very short. Dmochowski [17] pointed out the trade-off between intensity performance and focality performance in the MLS scheme. From our investigations (Fig 3 & 4), we found that changing the expected electrical field strength *e*_0_ affected this trade-off. Lower e0 can increase the focality of stimulation while higher e0 can increase the intensity. If intensity and focality performance are both important, it may be preferable to search the grid for the expected electrical field strength. However, this operation is known to be time consuming.

As an improvement to the MLS scheme, the ME scheme could achieve the maximum intensity performance that MLS scheme could be or might be reached (Fig 4) under power constraints, given in Eq 18. The advantage of the ME scheme is that it is only necessary to determine the expected stimulation direction, while for the MLS scheme, the expected stimulation strength *e*_0_ and direction must be determined. However, this advantage prevented the trade-off between intensity and focality from being controlled, leading to the best intensity performance but the worst focality performance when compared with the MLS scheme (Fig 4). Thus, in the single target case, we recommend the ME scheme for the best intensity performance, and for increased accuracy in controlling the intensity and focality performance, we recommend the MLS scheme.

From the Eq 8 and Eq 11, it is the definition that makes MLS and ME scheme have different performance in intensity and focality. The optimization problem of MLS scheme is to minimize the mean square error between simulation electrical field and the desired one. Thus, it can achieve the best focality performance. The optimization proble of ME scheme is to maximum the stimulation intensity of target ROI. Thus, it can achieve the best intensity performance.

### Multi-Target Case

Normal brain function requires cooperation among many brain regions. Thus, simultaneously applying stimulation to two or more brain regions may be more meaningful in both therapeutic and research applications. We found that directly using the MLS scheme or ME scheme (e.g., taking all target regions as a whole) for multi-target optimization caused the ‘overfitting’ problem, i.e., the optimization algorithm tended to optimize those regions that were larger in size or had a greater impact on the cost function, and ignore other regions (the first column in Fig 5 and the Fig 6). Thus, we used a weighted MLS scheme and weighted ME scheme to overcome this problem. Our results shown that when applying appropriate weights to each ROI, the weighted MLS scheme and weighted ME scheme could implement an arbitrary regional based stimulation mode (the second column to the fifth column in Fig 5 and Fig 7).

For the weighted ME scheme, besides applying weights to each ROI, additional regional strength distribution constraints or current distribution constraints should also be added due to the ‘maximum’ optimization mode. In this optimization mode, if one ROI has the largest effect on the cost function, distributing all current to that ROI will increase the cost function to the maximum value. If all ROIs have the same effect on the cost function, arbitrary current distribution will make the cost function to the maximum value. Thus, if additional distribution constraints are not added, the optimization results will maybe wrong. For the weighted MLS scheme, simply applying weights to each ROI is enough.

As with single target stimulation, the weighted ME scheme had the best intensity performance, but the stimulation distribution inside the ROI could not be controlled. Although the weighted MLS scheme could control the trade-off between intensity and focality when given an appropriate expected electrical field strength, searching for this value was somewhat time consuming.

### Multi-Target Under a Specific Task

To further validate the optimization ability of the weighted MLS scheme and weighted ME scheme in a multi-target case, we used a specific SWMT activation T-map as the stimulation mode.

For the weighted ME scheme, considering its regional based nature, we used a stimulation mode in which each ROI had a sum of stimulation strength proportional to its peak T value. Our results showed that when applying appropriate weights and additional strength distribution constraints or current distribution constraints, the weighted ME scheme could accurately distribute the expected stimulation strength to each ROI (Table 5). These results not only demonstrated the validity of our proposed weighted method, they also demonstrated the excellent stimulation energy distribution ability of the weighed ME scheme. However, as shown in Fig 9, when ROI 3 was relatively large and ROI 2 was relatively small, although the stimulation energy was precisely distributed to each ROI, the total sum of the stimulation energy was so small that some larger ROIs received almost no stimulation from visual aspect. Additionally, the sum of the electrical field strength was very small compared to the values in Table 3. Because ROI 2 was much smaller than L SFG in Table 3 (439 vs. 4458 voxels, about 1:10), the maximum sum of the electrical field strength was also very small, resulting in a reduction of the total brain stimulation level. Thus, when using the weighted ME scheme, a stimulation mode that incorporates stimulation energy distribution might be more suitable for situations in which all ROIs have similar sizes.

For the weighted MLS scheme, we used the SWMT activation T-map to directly apply weights to each voxel [21]. Our results showed that directly applying weights might also cause ‘overfitting’ if some of these weights are too large due to the substantial imbalance in ROI size. This problem could be dealt with to some extent by appropriately increasing the weights of the ROIs with small weights (Fig 10 and Table 7). However, from Table 6 we see that even though the overfitting problem decreased, the actual focality performance of the weighted MLS scheme was still worse. Thus, the weighted MLS scheme proposed in this study might be not suitable for this stimulation mode. Future research will focus on improving methods and searching for new optimization schemes for use with this stimulation mode. Although there needs to be improved, our proposed methods can still precisely distribute the energy into each target brain region according to any given ratio (Table 5). Thus, our methods can become a guideline for those studies or clinical treatments which need to stimulate two or more brain regions simultaneously using tDCS.

### Limitations

Firstly, in this study, an interior-point method was used to solve optimization problem. This method is a local search method under which the optimization result relies on the selection of initial point. In this study, we repeated solving the same optimization problem with a random initial point 100 times and chose the best result as the final solution, which can be very time consuming. Additionally, we used annealing and genetic algorithms to solve the optimization problem. However, we found that the optimization results of these two algorithms are similar with interior-points method, and more time will be cost by these two algorithms thanrunning the interior-points method 100 times. Thus, Future work will focus on the improvement of these two optimization schemes and finding more effective algorithms to solve optimization problems.

Secondly, as shown in Fig 3, stimulation depth is a problem for tDCS electrode displacement using the EEG 10/10 system. Grossman [?] proposed a novel beat frequency stimulation mode which had better focality and stimulation depth compared with dense electrode array tDCS. In future work, we plan to simulate this stimulation method using a realistic head model.

Finally, our method for constructing head models is not optimal. Furthermore, as reported previously [28,29], the more precise tissue segmentation and the value of conductivity *σ* for each tissue type need to be carefully considered. Thus, in future work, we plan to improve the construction methods and parameter settings of our head model.

## CONCLUSION

This study validates previous works in [17] and [18] in single target stimulation case using MLS and ME optimization schemes, then adds upon their work, by investigating the performance these two schemes in multi-target stimulation case. Our findings suggest that directly using MLS or ME scheme in multi-target stimulation case (e.g. taking all stimulation ROIs as a whole) may lead to a wrong stimulation result. By using our proposed weighted MLS and weighted ME schemes, this confusing result will be greatly improved. Thus, for any expected stimulation mode, carefully consideration of the weight of each stimulated ROIs is needed. Future work will focus on the improvements of our proposed methods.

## FUNDING

This work was financially supported by National Basic Research Program of China under Grant Nos. 2015CB856403 and 2014CB543203, the Engineering Research Center of Artificial Intelligence of Xi’an under Grant 201809170CX11JC12, and the National Natural Science Foundation of China under Grant Nos. 81471811 and 81471738.

## ACKNOWLEDGEMENT

We thank Lesley McCollum, PhD, from Liwen Bianji, Edanz Editing China (www.liwenbianji.cn/ac), for editing the English text of a draft of this manuscript.

## APPENDIX

We show the derivation of *α* in Fig 9. According to the specific stimulation parameters, Eq 25 can be written as:

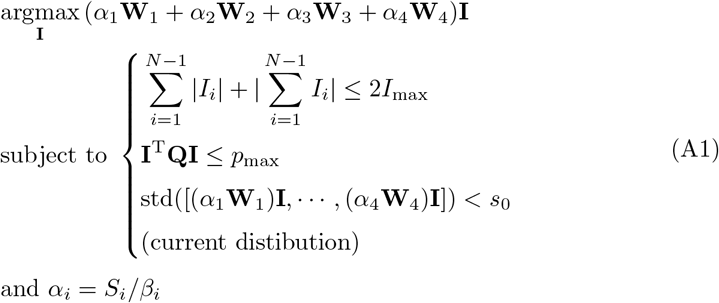

Where

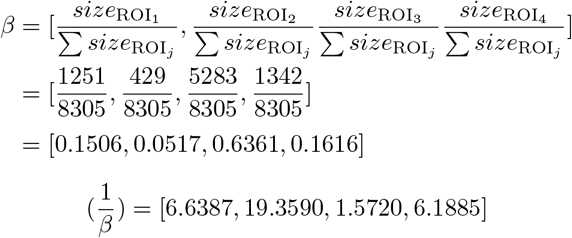

The desired stimulation strength ratio is,

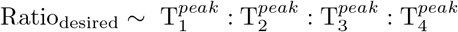

Thus, after current distribution, we get:

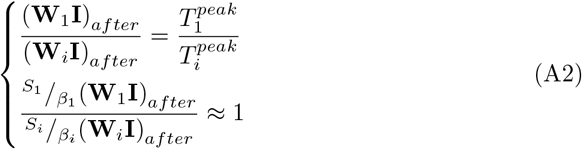

Solving the Eq A2, we get:

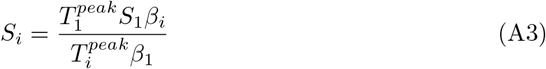

Let *S*_1_ = 1, we get:

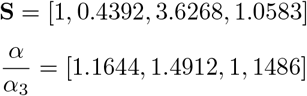

